# Changes in chromatin accessibility are not concordant with transcriptional changes for single-factor perturbations

**DOI:** 10.1101/2022.02.03.478981

**Authors:** Karun Kiani, Eric M. Sanford, Yogesh Goyal, Arjun Raj

**Affiliations:** Genetics and Epigenetics, Cell and Molecular Biology Graduate Group, Perelman School of Medicine, University of Pennsylvania, Philadelphia, PA, USA; Genomics and Computational Biology Graduate Group, Perelman School of Medicine, University of Pennsylvania, Philadelphia, PA, US; Department of Bioengineering, School of Engineering and Applied Sciences, University of Pennsylvania, Philadelphia, PA, USA; Department of Cell and Developmental Biology, Feinberg School of Medicine, Northwestern University, Chicago, IL, USA; Center for Synthetic Biology, Northwestern University, Chicago, IL, USA; Department of Genetics, Perelman School of Medicine, University of Pennsylvania, Philadelphia, PA, USA

## Abstract

A major goal in the field of transcriptional regulation is the mapping of changes in the binding of transcription factors to the resultant changes in gene expression. Recently, methods for measuring chromatin accessibility have enabled us to measure changes in accessibility across the genome, which are thought to correspond to transcription factor binding events. In concert with RNA-sequencing, these data in principle enable such mappings; however, few studies have looked at their concordance over short duration treatments with specific perturbations. Here, we used tandem, bulk ATAC-seq and RNA-seq measurements from MCF-7 breast carcinoma cells to systematically evaluate the concordance between changes in accessibility and changes in expression in response to retinoic acid and TGF-β. We found two classes of genes whose expression showed a significant change: those that showed some change in accessibility of nearby chromatin, and those that showed virtually no change despite strong changes in expression. The peaks associated with genes in the former group had a lower baseline accessibility prior to exposure to signal. Focusing the analysis specifically on peaks with motifs for transcription factors associated with retinoic acid and TGF-β signaling did not reduce the lack of correspondence. Analysis of paired chromatin accessibility and gene expression data from distinct paths along the hematopoietic differentiation trajectory showed a much stronger correspondence, suggesting that the multifactorial biological processes associated with differentiation may lead to changes in chromatin accessibility that reflect rather than drive altered transcriptional status. Together, these results show many gene expression changes can happen independent of changes in accessibility of local chromatin in the context of a single-factor perturbation and suggest that some changes to accessibility changes may occur after changes to expression, rather than before.

## Introduction

Transcription factors regulate gene expression by binding to specific DNA sequences, facilitating transcription through the recruitment and activation of the transcriptional machinery. Deciphering the combinatorial logic underlying which transcription factors bind to what portions of DNA and in what contexts is a central challenge in creating a complete model of transcriptional regulation. Sequencing-based methods have enabled the measurement of transcript levels for all genes as well as the putative binding profiles of transcription factors across the genome. However, the precise mapping between changes in these putative binding profiles and the changes in transcriptional activity remain the subject of debate.

A key component of decoding the relationship between transcription factor activity and the resultant changes in transcription is the measurement of transcription factor binding to DNA. Recently, the combination of biochemical binding assays with sequencing-based readouts has led to a cornucopia of methods for making such measurements. One workhorse method is chromatin immunoprecipitation sequencing (ChIP-seq), which characterizes the binding of transcription factors and other DNA-protein interactions genome-wide [1–3] by using immunoprecipitation of proteins that bind to chromatin and subsequently sequencing the coprecipitated DNA. However, ChIP-seq is limited in that each experiment can only interrogate the binding profile of one transcription factor at a time.

An alternative approach that circumvents that issue is the measurement of changes in accessibility of DNA to infer changes in the binding of all transcription factors at once. Accessible regions of DNA (i.e. those regions depleted of nucleosomes) represent only 3% of the genome, but often participate in the regulation of gene expression [4–7]. These regions can be detected genome-wide by combining the enzymatic activity of nucleases with high-throughput sequencing using techniques such as DNase I hypersensitive site sequencing (DNase-seq) [8] and assay for transposase accessible chromatin with sequencing (ATAC-seq) [9]. The interpretation of these accessibility methods leans heavily on the assumption that changes in regulatory factor binding are reflected in changes in chromatin accessibility. Certainly, there are many examples in which the correspondence between changes in accessibility strongly correspond to changes in transcriptional output. For instance, summation of ChIP-seq signal for 42 transcription factors mapped by encode in K562 chronic myelogenous leukemia cells paralleled the signal from accessible sites revealed by DNase-seq [7]. Moreover, computational methods to infer transcription factor footprints from accessibility measurements have been shown to recapitulate ChIP-seq binding well [10]. Accessibility methods can also be used to look for changes in accessibility across various perturbations and cell types. Changes in accessibility generally seem to correspond to changes in transcription in the sense that large changes in transcriptional output are reflected in broad changes in the accessibility of several loci in the surrounding chromatin [11,12].

However, it is unclear how well these accessibility based methods capture the activity of all transcription factors. It is possible that some transcription factors’ binding and activity does not result in corresponding changes in accessibility and vice versa. Such a lack of correspondence could manifest itself as a lack of correlation between changes in accessibility and changes in transcription. Given the underlying assumption that a change in transcription must be mediated by the change in some transcription factor activity, then such a lack of correspondence would suggest that changes in the activity of transcription factors could change expression without changing accessibility near its binding site. While reports from the literature generally show a strong correspondence [11–14], it is worth noting that the comparisons in such studies are often across rather different cell types. In such cases, it is possible that the changes in accessibility are not driven by regulation *per se*, but rather reflect the consequences of sequential exposure to multiple regulatory factors that characterize the differentiation process. Such accessibility changes could, in principle, signify the reinforcement of genes that are already transcriptionally active genes, or could even just appear around actively transcribed genes without any functional role. Disentangling such possibilities could be revealed with the use of single-factor perturbations that more directly affect an individual pathway; however, few such data are available.

Here, we used tandem bulk RNA-seq and ATAC-seq data from MCF-7 breast carcinoma cells exposed to multiple doses of retinoic acid or TGF-β to determine the degree of concordance between changes in chromatin accessibility and changes in gene expression. Furthermore, we evaluated concordance in another published data set of hematopoietic differentiation to validate our approach based on well-defined and specific perturbations. We demonstrate that while some differentially expressed genes have a high concordance between gene expression and chromatin accessibility changes, many other genes are differentially expressed without changes in their local chromatin accessibility.

## Results

### Genome-wide expression and chromatin accessibility changes reflect known biology of two perturbations

To measure the correspondence between changes in chromatin accessibility and changes in gene expression, we used MCF-7 breast carcinoma cells due to their previously described transcriptional responses to all-*trans* retinoic acid [15] (referred to from here on as retinoic acid) and transforming growth factor beta (TGF-β) [16]. We used paired, bulk accessibility (ATAC-seq) and expression data (RNA-seq) from these cells [17] collected 72 hours after continuous exposure to three different doses of each signal (Figure 1A). We chose this timescale because previous work with MCF-7 cells showed more transcriptional changes at 72 hours compared to 24 hours after exposure to retinoic acid [15], and chromatin accessibility changes may not be detectable until 24 hours after perturbation [18]. Differential gene expression and differential peak accessibility analysis showed a dose-dependent response to both signals compared to ethanol control (Figure 1A, bar plots). The ethanol ‘vehicle’ controls comprise three different densities of cells, and the transcriptomes of control conditions globally were similar regardless of cell density (Supplemental Figure 1A). To confirm that global gene expression and chromatin accessibility patterns were similar between signals and dosages, we performed principal component analysis. For both RNA-seq and ATAC-seq data, all samples exposed to the same signal or ethanol control clustered together, indicating that their gene expression and chromatin accessibility were more similar to each other than to other conditions, supporting the quality of these data.

**Figure 1:**
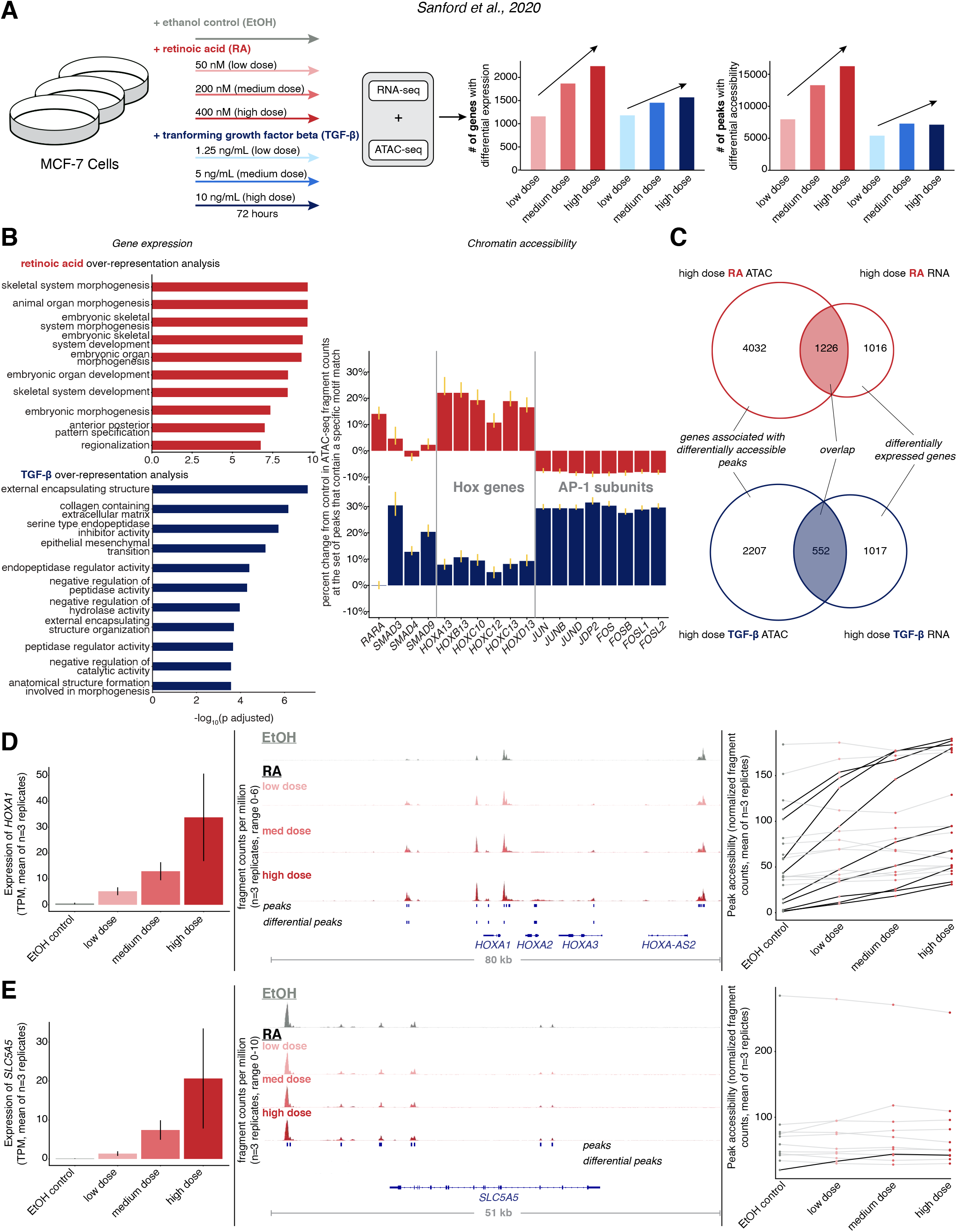
Changes in gene expression can occur with or without concordant changes in chromatin accessibility in response to signal. a. Schematic of signal response experiments in MCF-7 cells from Sanford et al., 2020. Briefly, cells were treated with either ethanol vehicle control (gray) or three different doses of retinoic acid (shades of red) or TGF-β (shades of blue). After 72 hours of continuous exposure, bulk RNA-seq and ATAC-seq were performed on samples. We show the number of differentially expressed genes and differentially accessible peaks for each dose of each condition compared to ethanol vehicle control. b. Validation that changes in gene expression and chromatin accessibility reflect known biology of perturbations. Left: overrepresentation analysis of differentially upregulated genes in response to high dose retinoic acid (red) or TGF-β (blue). Top ten gene sets for each signal by −log_10_ FDR-adjusted p-value are shown. Right: motif enrichment analysis of differentially accessible peaks for selected motifs of transcription factors related signaling pathways of these signals. Y-axis shows percentage change of ATAC-seq signal at motif containing peaks relative to ethanol vehicle control samples. For each condition, we pooled together replicates for all three doses. Error bars represent bootstrapped confidence intervals. c. Overlap between changes in gene expression and changes in chromatin accessibility in response to high dose retinoic acid (top) or high dose TGF-β (bottom). Of the genes that were differentially expressed (right circle of Venn diagram) we looked at the overlap (shaded) of how many of them also had at least one differentially accessible peak (left circle). To disprove the null hypothesis that there is no association between genes that are differentially expressed and genes that have differentially accessible peaks assigned to them using the ‘nearest’ approach, we performed Fisher’s exact test to show the probability of these data or more extreme if the null hypothesis was true for both signals was less than 2.2×10^-16^. d. Expression and accessibility change of *HOXA1* in response to increasing doses of retinoic acid. Left: Expression (TPM, triplicate average) in response to increasing dose of retinoic acid (error bars represent SEM). Middle: track view of *HOXA1* locus with accessibility in fragments per million and peaks and differential peaks annotated. Right: quantification of peak accessibility (normalized fragment counts, triplicate average) within a 50 kilobase window of *HOXA1* locus with peaks that are differentially accessible between ethanol vehicle control and high dose retinoic acid conditions marked with black lines. e. Expression and accessibility change of *SLC5A5* in response to increasing doses of retinoic acid. Left: Expression (TPM, triplicate average) in response to increasing dose of retinoic acid (error bars represent SEM). Middle: track view of *SLC5A5* locus with accessibility in fragments per million and peaks and differential peaks annotated. Right: quantification of peak accessibility (normalized fragment counts, triplicate average) within a 50 kilobase window of *SLC5A5* locus with peaks that are differentially accessible between ethanol vehicle control and high dose retinoic acid conditions marked with black lines.

To validate that changes in gene expression were consistent with the known biology of these signaling pathways, we performed over-representation analysis on the upregulated genes in response to high dose retinoic acid or TGF-β against curated gene sets from the molecular signatures database [19,20]. The top ten gene sets based on false discovery rate (FDR)-adjusted p-values were processes canonically associated with retinoic acid (*morphogenesis*, *organ development*, *anterior-posterior patterning*) and TGF-β (*extracellular matrix*, *endopeptidase activity*), respectively (Figure 1B). Gene set enrichment analysis [21] showed that genes that were differentially expressed in response to high dose retinoic acid were significantly enriched for genes associated with skeletal system morphogenesis, and genes that were differentially expressed as a result of exposure to high dose TGF-β were significantly enriched for genes associated with epithelial-to-mesenchymal transition (Supplemental Figure 1B). Thus, the differentially expressed genes generally reflected the known biology of the signals the cells were exposed to.

We next wondered if the changes in chromatin accessibility in response to signal were associated with the activity of specific transcription factors, in particular, those associated with the biology of these signaling pathways. We used a modified version of the chromVAR package along with its curated database of transcription factor motifs, cisBP, to identify the transcription factors with the largest predicted change in activity [22]. We used the set of differential peaks to determine the set of the top 150 transcription factors with the greatest magnitude of change. These included the binding motifs of transcription factors that are canonical effectors of retinoic acid (RAR-α, HOXA13) and TGF-β signaling (SMAD3, SMAD4, and SMAD9). For each of these transcription factor motifs, we calculated a motif enrichment score for each condition based on the bias-uncorrected deviation score from chromVAR. The motif enrichment score represents the percentage change in ATAC-seq fragment counts in all peaks that contain a given transcription factor’s motif (Figure 1B). For example, the enrichment score of 28% for SMAD3 in the TGF-β condition meant that peaks containing the SMAD3 motif on average saw a 28% increase in fragment counts after exposure to TGF-β. We pooled together the low, medium, and high doses for each condition together in order to decrease the variability of motif enrichment scores estimates. Thus, our data recapitulated expected changes in accessibility, presumably due to the activity of transcription factors well-known to be activated by the signals used. Thus, of the changes in accessibility we did detect, they made sense based on a model of transcription factor activity leading to changes in accessibility. However, it was still possible that the activity of many transcription factors was not captured by changes in accessibility.

### The relationship between changes in chromatin accessibility and gene expression varies on a gene by gene basis

We next wondered whether genes that were differentially expressed were more likely to have differentially accessible peaks nearby, i.e., was there concordance between gene expression and chromatin accessibility changes at the level of individual genes? To characterize the extent of concordance between these data, we looked at the overlap between genes that were differentially expressed in response to high dose signal and genes with differentially accessible peaks nearby after exposure to signal (Figure 1C). We assigned each accessible peak to the nearest transcriptional start site (“nearest approach”) and found that of the over 2000 genes upregulated in response to high dose retinoic acid, more than half of them had at least one differential peak assigned to its transcriptional start site (p-value < 2.2×10^-16^, Fisher’s exact test). Similarly, a third of the genes whose expression was upregulated in response to TGF-β had differential peaks assigned to them (p-value < 2.2×10^-16^, Fisher’s exact test). Thus, genes that are differentially expressed are more likely than random chance to have a nearby peak that is differentially accessible in response to retinoic acid or TGF-β.

While using this overlap-based approach showed correspondence between genes that are differentially expressed and their nearby peaks in response to signal, aspects of the nature of the concordance of these changes were not captured by this analysis. For example, the overlap-based method counted all differentially accessible genes that had at least one differentially accessible peak assigned to them as concordant, but did not take into account the proportion or degree to which those nearby peaks change. Moreover, we did not take into account the relationship between directionality of changes in gene expression and chromatin accessibility. The underlying assumption at the basis of this relationship is that when peaks become more accessible that the nearby gene increases its expression, and the overlap-based approach does not take this correspondence of the direction of change into account. To better characterize these facets of concordance, we first individually examined the changes in chromatin accessibility nearby two genes whose expression were upregulated in response to retinoic acid.

*HOXA1* and *SLC5A5* induction are associated with exposure to retinoic acid [23–25], and both genes showed a dose-dependent increase in expression in response to retinoic acid (Figures 1D,E leftmost panels). After optimizing parameters for calling peaks and determining differentially accessible peaks (Supplemental Figure 2), we found that while a large number of peaks are differentially accessible near the *HOXA1* locus (Figure 1D, track view middle, black traces in accessibility plot, right), very few peaks are differentially accessible near the *SLC5A5* locus (Figure 1E, track view middle, accessibility plot, right). Therefore, genes with high expression change in response to signal can show a large degree of accessibility changes or show very little accessibility changes, suggesting that changes in transcription factor activity may or may not be reflected in changes in accessibility.

### Chromatin accessibility changes are less concordant with large changes in gene expression in signaling compared to hematopoietic differentiation

Next, we evaluated the concordance between accessibility and gene expression genome-wide while also factoring in the directionality of changes and the relative proportion of peaks that are changing on a gene by gene basis. As a point of comparison, we used previously published gene expression and chromatin accessibility data from hematopoietic differentiation [11] that demonstrated that large changes in gene expression were typically associated with gains or losses (depending on the direction of expression change) of cell type-specific enhancers when comparing the expression and accessibility of hematopoietic stem and progenitor cells (HSPCs) to monocytes.

Before using this data set as a comparison to ours for measuring concordance between chromatin accessibility and gene expression changes, we verified that the hematopoietic differentiation data was similar to our own by a variety of metrics. First, we wanted to compare whether the number of differentially expressed genes and differentially accessible peaks between HSPCs and monocytes in the hematopoietic differentiation data was similar to the numbers from MCF-7 cells exposed to retinoic acid or TGF-β. We found that both HSPC and monocyte populations had greater than 2000 genes that were specifically expressed in their respective cell types compared to the approximately 2000 and 1500 genes differentially expressed in MCF-7 cells in response to high dose retinoic acid and TGF-β, respectively (Figure 1A). Moreover, HSPC and monocyte populations had more than 6000 differentially accessible peaks (Supplemental Figure 3A) compared to the approximately 15000 and 6000 differentially accessible peaks in MCF-7 cells in response to high dose retinoic acid and TGF-β, respectively (Figure 1A).

Next, we annotated the location of peaks based on where in the genome they were located relative to gene bodies and quantified what proportion of peaks fell into annotation categories such as promoter, intergenic, exonic, intronic, etc. ATAC-seq peaks from MCF-7 cells had a larger proportion of peaks at gene promoters (within 3 kilobases upstream or downstream of the transcription start site) whereas a greater proportion of the DNase I hypersensitive sites in the HSPC and monocyte populations were from distal intergenic regions compared to promoters (Supplemental Figure 3B). This finding could be the result of inherent differences in the assays or could reflect biological differences. Moreover, the MCF-7 data had a greater proportion of peaks located at gene promoters, which could in principle bias our results toward having a larger degree of concordance because accessibility changes at promoters were more strongly correlated with gene expression changes than distal accessible. Despite this bias, our data demonstrate less concordance.

Given the different assays used to determine genome-wide chromatin accessibility, we realigned the DNase-seq data to the hg38 reference and examined the peaks at a ‘housekeeping gene’ (*GAPDH*), hematopoietic differentiation-specific genes (*CD34*, *CD14*) and retinoic acid and TGF-β-related genes (*DHRS3*, *SERPINA11*) to spot-check that the accessibility data were similar. Indeed, there were similar accessibility profiles for *GAPDH*, and appropriate differences in accessibility given the cell type of signal for the other sites, indicating the accessibility data were comparable (Supplemental Figure 4A-E). Moreover, to look at similarities in accessibility genome-wide, we calculated the intersection of the consensus peak sets from hematopoietic differentiation and MCF-7 signal response data sets. We observed that approximately 55% of peaks from hematopoietic differentiation data (DNase-seq) overlapped with peaks from the MCF-7 signal response data set (ATAC-seq). These results show that the datasets do not have systematic qualitative differences in either expression or accessibility, enabling us to compare the degree of concordance across these two systems.

In the original analysis of hematopoietic differentiation, the authors found that regulatory complexity (defined as the number of accessible regions closest to a gene’s transcriptional unit) was an important discriminating factor for whether changes in accessibility corresponded to changes in expression, with areas of high complexity showing more correspondence than those of low complexity. Hence, we similarly grouped genes from our MCF-7 dataset into high and low complexity for our comparisons. We categorized genes with more than 7 peaks assigned to them using the ‘nearest approach’ as ‘high complexity’, while genes with 7 or fewer peaks were categorized as having ‘low complexity’ (Figure 2A, top panel). The cutoff for loci complexity was calculated by taking a tertile based approach [11] and calling any number of peaks above the highest tertile cutoff as high and any peak below that as low complexity (Figure 2B, solid line, lower plot). Because high complexity genes on average had higher levels of expression in the hematopoietic differentiation data, we sought to determine if there was any difference in expression between high and low complexity genes in our MCF-7 data. The median expression of high complexity loci was similarly higher than low complexity loci in response to both exposure to high dose retinoic acid (23.30 versus 13.27 TPM) and high dose TGF-β (24.06 versus 13.05 TPM) (Supplemental Figure 5A, p-value < 2.2×10^-16^ for both, Kolmogorov-Smirnov test) demonstrating that high complexity genes are more highly expressed as in the hematopoietic differentiation data. Despite this difference in expression, the distributions of peak widths for peaks of high and low complexity genes were similar (Supplemental Figure 5B).

**Figure 2:**
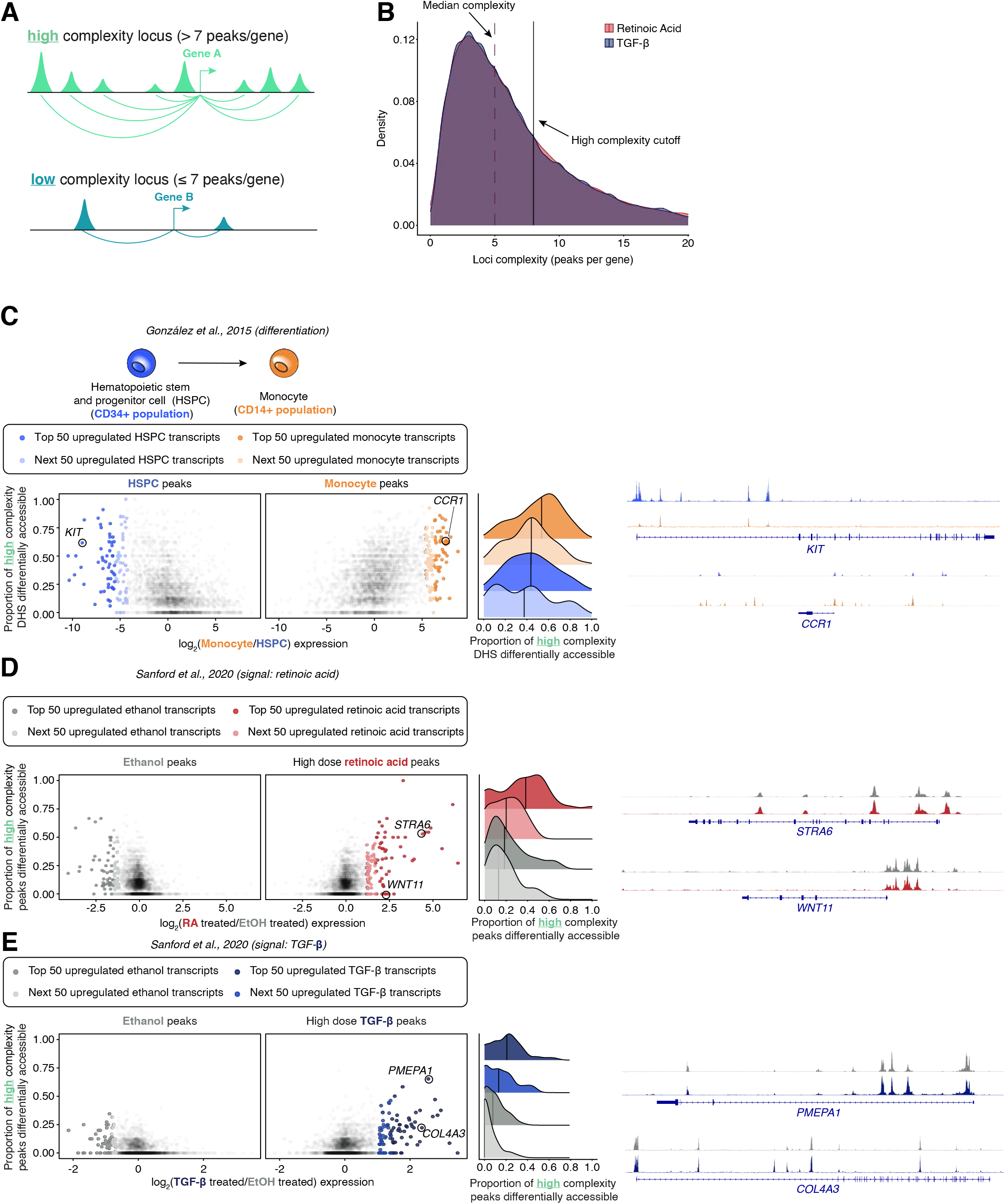
Signaling shows less concordance between highly differentially expressed genes and chromatin accessibility changes compared to hematopoietic differentiation data for high complexity genes. a. Schematic demonstrating classification of genes into “high” versus “low” complexity genes based on the number peaks assigned to a gene using the ‘nearest’ approach. b. Density plot of number of peaks per gene in retinoic acid (red) and TGF-β (blue, overlap in purple) with median complexity marked by dotted line and high complexity cutoff marked by solid line. c. Concordance between expression and accessibility changes between hematopoietic stem and progenitor cells and monocytes. Left: plot showing changes in gene expression in CD34+ hematopoietic stem and progenitor cells (blue) and CD14+ monocytes (orange) from González et al., 2015 (schematic, top). For the plots, each dot is a gene, and on the x axis is log_2_fold change in expression and on the y-axis the proportion of differentially accessible DHSs for each associated gene. The top 100 most highly expressed genes in hematopoietic stem and progenitor cells and monocytes are colored in shades of orange and blue, respectively. Middle: density plot of the distribution of the proportion of high complexity DHS associated with the top 100 expressed genes in CD34+ hematopoietic stem and progenitor cells and CD14+ monocytes with median value marked by vertical black line. Right: example tracks DNase I sequencing data for *KIT* and *CCR1* (marked on plot on left). d. Concordance between expression and accessibility changes between cells exposed to ethanol vehicle control and high dose retinoic acid. Left: plot showing changes in gene expression and chromatin accessibility between ethanol vehicle control and high dose retinoic acid. Each dot is a gene, and on the x axis is the log_2_ fold change in expression and on the y-axis the proportion of differentially accessible ATAC-seq peaks for each gene. The top 100 most highly expressed genes in ethanol vehicle control and high dose retinoic acid are colored in shades of gray and red, respectively. Middle: density plot of the distribution of the proportion of high complexity ATAC-seq peaks associated with the top 100 expressed genes in ethanol vehicle control and high dose retinoic acid with median value marked by vertical black line. Right: example ATAC-seq tracks of *STRA6* and *WNT11*. e. Concordance between expression and accessibility changes between cells exposed to ethanol vehicle control and high dose TGF-β. Left: plot showing changes in gene expression and chromatin accessibility between ethanol vehicle control and high dose TGF-β. Each dot is a gene, and on the x axis is the log_2_ fold change in expression and on the y-axis the proportion of differentially accessible ATAC-seq peaks for each gene. The top 100 most highly expressed genes in ethanol vehicle control and high dose TGF-β are colored in shades of gray and blue, respectively. Middle: density plot of the distribution of the proportion of high complexity ATAC-seq peaks associated with the top 100 expressed genes in ethanol vehicle control and high dose retinoic acid with median value marked by vertical black line. Right: example ATAC-seq tracks of *PMEPA1* and *COL4A3*.

We began our analysis by focusing on the high complexity genes. To determine the concordance between gene expression changes and chromatin accessibility changes, we used the ‘nearest approach’ to assign peaks to genes. For each gene we compared the log_2_ of the fold change in expression between conditions versus the proportion of peaks that were differentially accessible in the same direction (i.e., peaks that increase in accessibility for genes that increase in expression after exposure to signal and vice versa). We observed that for hematopoietic differentiation, the 100 most highly expressed high complexity genes in the HSPC and monocyte populations had a high proportion of peaks which were differentially accessible in the concordant direction, reproducing the conclusions of González et al. that large changes in expression were consistently associated with concordant changes in chromatin accessibility (Figure 2C). Next, we used this approach on our data to compare expression and accessibility changes between ethanol vehicle control and high dose retinoic acid or TGF-β. For both signals, we observed two distinct groups of genes within the top 100 most differentially expressed genes. One group of genes (‘accessibility-concordant genes’) behaved similarly to those in the hematopoietic differentiation data, demonstrating a concordance between expression and accessibility changes (Figures 2D,E). However, the other group of genes (‘accessibility-non-concordant genes’) had large expression changes with little to no peaks nearby changing in accessibility, creating a skew in the distribution toward a lower proportion of peaks being differentially accessible in a concordant manner compared to the hematopoietic differentiation data (Figures 2C-E, density plots).

Adjusting the minimum peak coverage parameter changes the number of differential peaks and the proportion of differential peaks that change in the corresponding direction of expression. We wondered if a lower minimum coverage threshold changed the qualitative result we noticed before and thus conducted the same analysis using a lower minimum peak coverage threshold for determining differential peaks (see methods). We observed that a similar pattern occurred in high complexity genes with this set of parameters (Supplemental Figure 6).

González and colleagues showed that for some low complexity genes, large changes in expression were not accompanied with concordant changes in accessibility [11]. We similarly wanted to confirm whether this decreased correspondence was the case in our data in response to retinoic acid and TGF-β. Using the same approach as before, we compared the log_2_ of the fold change in expression of low complexity genes to the proportion of peaks with differential accessibility in the concordant direction. The hematopoietic differentiation and signaling data for low complexity all qualitatively had genes whose expression increased without concordant changes in accessibility (Supplemental Figure 7A-C). The distribution of the proportion peaks that were differentially accessible in the concordant direction for the top 100 up and downregulated genes was roughly uniform when comparing HSPCs to monocytes (Supplemental Figure 7A, density plot on right). By comparison,the distribution was skewed toward more genes having a lower proportion of peaks being differentially accessible in the concordant direction in response to signals in MCF-7 cells, especially in the case of TGF-β (Supplemental Figure 7B,C, density plots on right). Thus, while both the signaling in MCF-7 and hematopoietic data demonstrated large gene expression changes without concordant changes in chromatin accessibility with low complexity genes, a greater proportion of genes did so in the signaling data.

### Peaks nearby genes with high concordance have lower accessibility prior to exposure to signal

We wondered what the differences were between genes that were differentially expressed and had large accessibility changes versus those that were differentially expressed and had low accessibility changes. First, for high dose retinoic acid and TGF-β, we split genes into four groups based on whether they were differentially expressed and the proportion of peaks assigned to them using the ‘nearest’ method that were differentially accessible in the appropriate direction. These four groups were (1) genes with differentially upregulated expression and concordant accessibility changes (2) genes with differentially upregulated expression non-concordant accessibility changes (3) genes with differentially downregulated expression and a concordant accessibility changes, and (4) genes with with differentially downregulated expression and non-concordant accessibility changes (Figures 3A,B). We quantified the distribution of peak complexity across these groups and observed that they were similar across all four gene subgroups (Supplemental Figures 8A,B).

**Figure 3:**
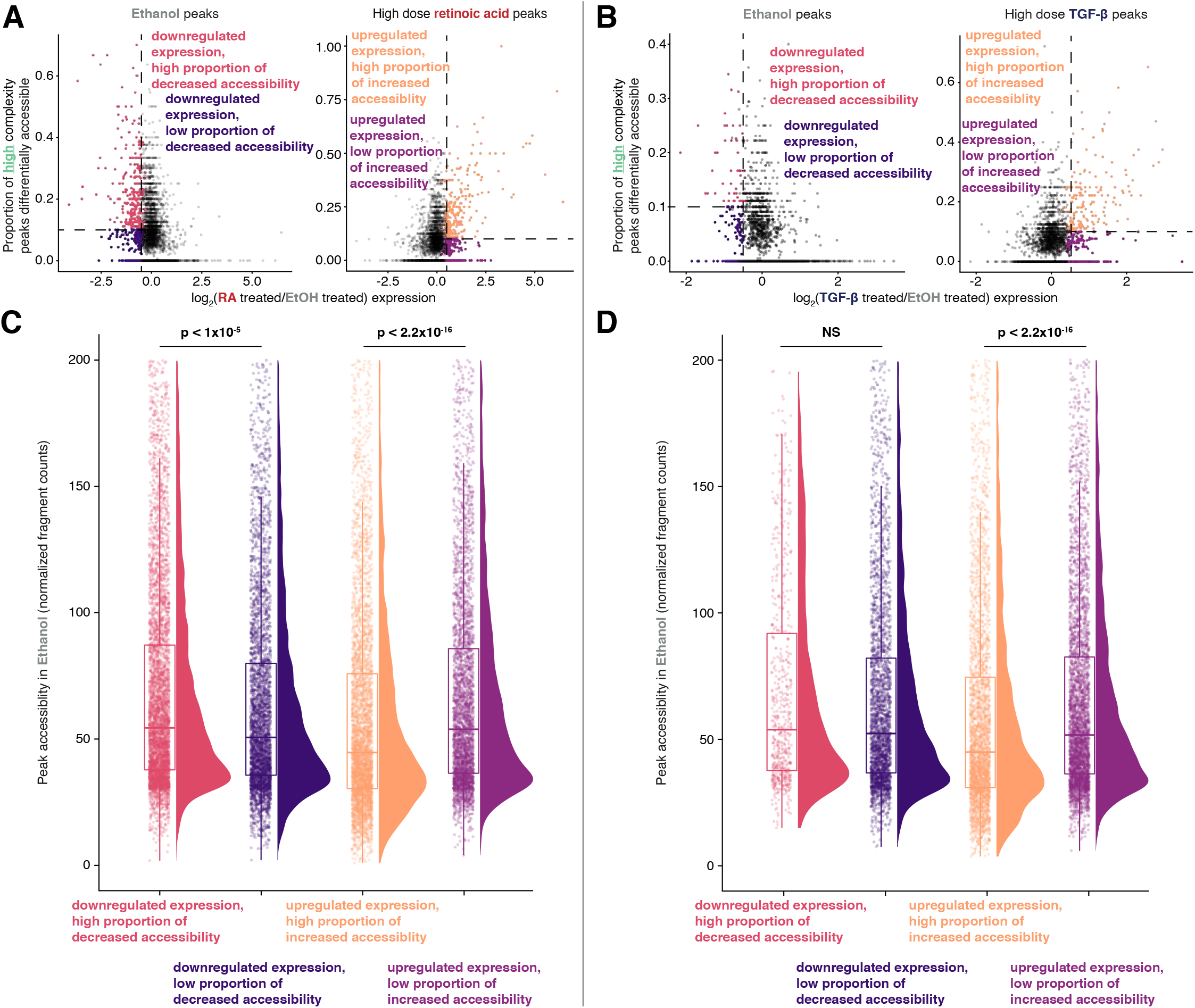
Separation of differentially expressed genes in response to signal into high and low concordance groups shows differences in pre-existing accessibility. a. Categorization of differentially expressed genes in response to high dose retinoic acid based on direction of expression change and proportion of peaks differentially accessible in the same direction. b. Categorization of differentially expressed genes in response to high dose TGF-β based on direction of expression change and proportion of peaks differentially accessible in the same direction. c. Differential accessibility in ethanol vehicle control conditions prior to addition of high dose retinoic acid. Accessibility of every peak assigned using the ‘nearest’ approach for gene groups from **(a)** in ethanol vehicle control conditions. P-values represent the probability of these data or more extreme under the null hypothesis that the distribution of peak accessibilities were drawn from the same probability distribution via the Kolmogorov-Smirnov test. d. Differential accessibility in ethanol vehicle control conditions prior to addition of high dose TGF-β. Accessibility of every peak assigned using the ‘nearest’ approach for gene groups from **(b)** in ethanol vehicle control conditions. P-values represent the probability of these data or more extreme under the null hypothesis that the distribution of peak accessibilities were drawn from the same probability distribution via the Kolmogorov-Smirnov test.

We first asked whether the change in accessibility between these two gene groups was due to differences in the preexisting accessibility of peaks for these genes. Indeed, we found the baseline accessibility of peaks for genes with concordant increases in expression and accessibility in ethanol vehicle conditions was lower than those of peaks of genes that increase in expression without a commensurate change in chromatin accessibility (Figure 3C). This relationship was also recapitulated for concordant peaks that increase in expression and accessibility in response to high dose TGF-β (Figure 3D). Similarly, when comparing genes that are differentially downregulated in expression a similar pattern holds true in the opposite direction (Figures 3C,D, Supplemental Figures 8C,D). One explanation may be that genes whose nearby chromatin was already accessible were permissive toward the action of the appropriate transcription factors to modulate expression. An alternative explanation is that the ATAC-seq assay itself had saturated in its ability to measure chromatin accessibility. In contrast, the difference in accessibility decreased between genes with a low proportion of peaks that were differentially accessible and genes with a high proportion of accessible peaks after exposure to signal Supplemental Figures 8C,D). Thus, the difference in the proportion of accessible peaks nearby the two groups of genes was partially explained by the pre-existing chromatin accessibility.

### Multiple approaches to integrating chromatin accessibility and gene expression changes show a low degree of concordance during signaling

Finally, we measured to what degree the change in accessibility of chromatin nearby a gene is reflected in the change in gene expression. Because linear distance is not always a good predictor of what accessible regions interact with what genes, we used multiple approaches to assign peaks to genes. First, we used the ‘nearest approach’ to create a one-to-one mapping between accessible sites and genes by assigning them to the nearest transcriptional start site [26,27], again comparing our signaling dataset to the hematopoietic differentiation dataset. Because many genes have multiple peaks assigned to them, we used two methods for collapsing peak values per gene: either the median accessibility of peaks across genes or the maximum (Figure 4A, schematic). We observed a stronger correlation between accessibility and expression changes in differentiation data (median approach Pearson’s r = 0.34, maximum approach Pearson’s r = 0.26) than in MCF-7 in response to signal (retinoic acid: median approach Pearson’s r = 0.27, maximum approach Pearson’s r = 0.10; TGF-β: median approach Pearson’s r = 0.27, maximum approach Pearson’s r = 0.10; Figure 4A, right side).

**Figure 4:**
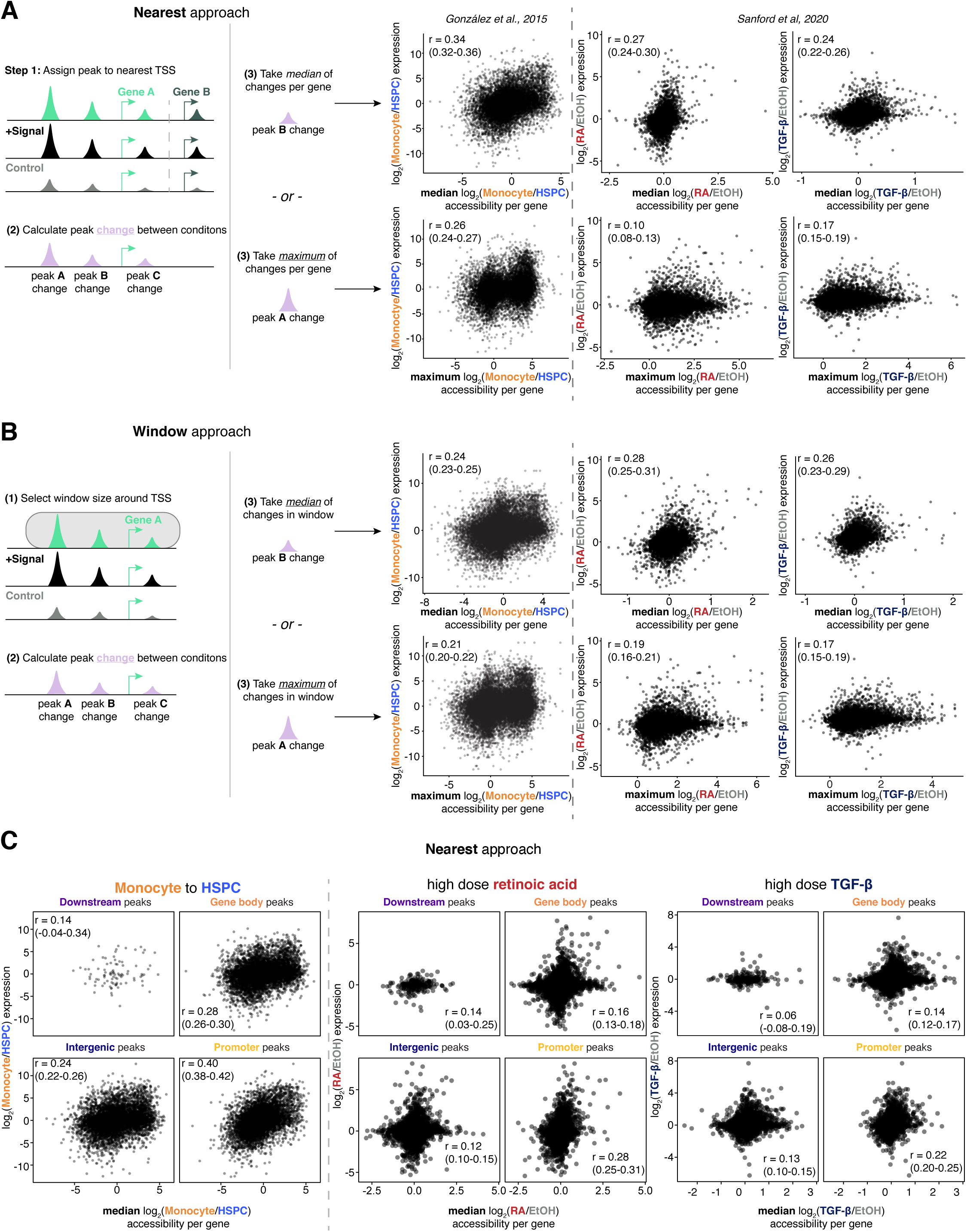
Multiple approaches to quantifying peak accessibility shows low correlation between gene expression changes and accessibility changes in signaling. a. ‘Nearest’ approach to assigning peaks to genes shows less concordance in signaling compared to hematopoietic differentiation. Left: schematic showing ‘nearest’ approach where peaks are assigned to the nearest transcriptional site and change in accessibility (purple) on a per-gene basis is calculated by either median change in accessibility (top row) or maximum peak change (bottom row). Right: scatter plots showing change in peak accessibility (median or maximum) versus log_2_ fold change in expression on y axis for hematopoietic differentiation data from González et al. (left column) and for high dose retinoic acid and high dose TGF-β (right two columns). Pearson’s correlation coefficients reported with 95% confidence interval from bootstrapping with 10,000 replicates in parentheses. b. ‘Window’ approach to assigning peaks to genes shows less concordance in signaling compared to hematopoietic differentiation. Left: schematic showing ‘window’ approach where all peaks within a certain window of the the transcriptional start site are assigned to that gene and the change in accessibility (purple) on a per-gene basis is calculated by the median change in accessibility (top row) or the maximum change in accessibility (bottom row). Right: scatter plots showing change in peak accessibility (median or maximum) using ‘window’ approach with a 100 kilobase window versus log_2_ fold change in expression on y axis for hematopoietic differentiation data from González et al. (left column) and for high dose retinoic acid and high dose TGF-β (right two columns). Pearson’s correlation coefficients reported with 95% confidence interval from bootstrapping with 10,000 replicates in parentheses. c. Using ‘nearest’ approach to look for correlation between accessibility and gene expression changes based on annotations of peak location. First two columns showing correlation for hematopoietic differentiation data from González et al, and right four columns showing correlation for high dose retinoic acid and high dose TGF-β, respectively. Pearson’s correlation coefficients reported with 95% confidence interval from bootstrapping with 10,000 replicates in parentheses.

Next, we used a window-based approach where there was the possibility of a many-to-one mapping of peaks to genes. We assigned all peaks within a 100 kilobase window [17] in order to maximize the number of differential peaks assigned to a gene (Supplemental Figure 9A,B). Similar to the ‘nearest’ approach, we collapsed values using median accessibility change across all peaks assigned to a gene as well as maximum accessibility per gene (Figure 4B, schematic) We observed a similar effect using this approach where there was a stronger correlation between change in accessibility and change in expression between HSPC versus monocyte versus MCF-7 cells exposed to signal (Figure 4B). Of note, the correlation coefficients were similar between both methods of assigning peaks.

We also wondered if the correlation between the extent of chromatin accessibility changes and gene expression changes would be different at the two lower doses. We used both the median and maximum peak value per gene while assigning peaks to genes using the nearest and window approaches. We observed similarly weak correlation as high dose signal using all methods at both low and medium doses (Supplemental Figure 9C,D). Consequently, the correlation between the magnitude of change in gene expression and chromatin accessibility was modest across the range of doses of signals.

To see if peaks in specific genomic regions (promoters, parts of the gene body, downstream and intergenic areas) had unique relationships between change in chromatin accessibility and change in gene expression, we subsetted our correlation analysis. We annotated peaks using ChiPseeker [28] to categorize them as being at promoters, within the gene body (5’ UTR, 3’ UTR, intronic, and exonic sequences), downstream of the gene end, or at intergenic sequences. We used peaks assigned to genes using the ‘nearest’ approach and took the median change in accessibility per gene. The strongest correlation between changes in accessibility and gene expression across sets of comparisons was at promoter peaks (Figure 4C). While promoter correlation is quantitatively stronger, the overall qualitative conclusion remains the same. Thus, despite using a variety of approaches for both assigning peaks to genes as well as collapsing the accessibility of all peaks for a given gene to a single value, we failed to appreciate a strong relationship between changes in accessibility and changes in gene expression.

Finally, we wondered if peaks that contained the motifs of transcription factors that are associated with retinoic acid and TGF-β signaling only (as opposed to all peaks) would show a stronger correlation between the changes in chromatin accessibility and gene expression. We annotated peaks with a log-likelihood score of a given motif being found in that peak and subsetted on those peaks with a nonzero log-likelihood score to examine the correlation between changes in accessibility and gene expression. Using this approach, we examined log-likelihood scores for motifs associated with retinoic acid signaling (RARA-α, HOXA13, and FOXA1) and motifs associated with TGF-β (SMAD3, SMAD4, and SMAD9). We observed that focusing on peaks annotated with peaks we would *a priori* expect to be involved in modulating gene expression in response to signal showed limited correlation between changes in chromatin accessibility and changes in gene expression (Figure 5).

**Figure 5:**
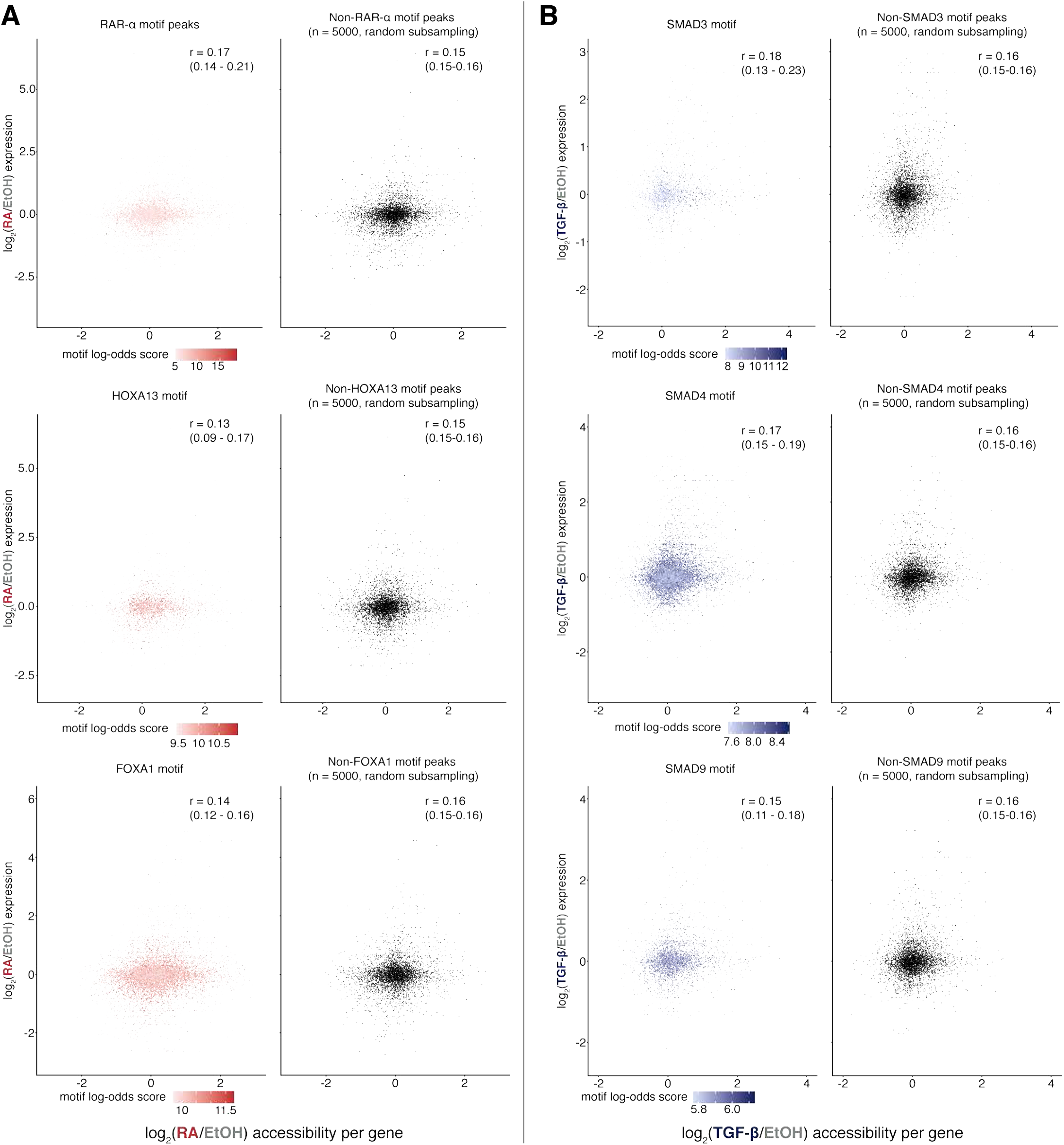
Focusing on peaks annotated for biologically relevant transcription factor motifs fails to demonstrate a strong correlation between the magnitude of gene expression and chromatin accessibility changes. a. Peaks annotated for motifs of transcription factors related to retinoic acid biology (RAR-α, HOXA13, FOXA1, left column) showed weak correlation between changes in gene expression and chromatin accessibility in response to high dose retinoic acid. Peaks are colored based on the log-odds of a motif being present in a given peak. Plot of expression and accessibility change for 5000 randomly sampled peaks lacking the corresponding peak (right column). Pearson’s correlation for peaks not having a given motif are for all peaks without that motif, not the 5000 subsampled peaks. Pearson’s correlation coefficients reported with 95% confidence interval from bootstrapping with 10,000 replicates in parentheses. b. Peaks annotated for motifs of transcription factors related to retinoic acid biology (SMAD3, SMAD4, SMAD9, left column) showed weak correlation between changes in gene expression and chromatin accessibility in response to high dose TGF-β. Peaks are colored based on the log-odds of a motif being present in a given peak. Plot of expression and accessibility change for 5000 randomly sampled peaks lacking the corresponding peak (right column). Pearson’s correlation for peaks not having a given motif are for all peaks without that motif, not the 5000 subsampled peaks. Pearson’s correlation coefficients reported with 95% confidence interval from bootstrapping with 10,000 replicates in parentheses.

## Discussion

Here, we integrated tandem, genome-wide chromatin accessibility and transcriptomic data to characterize the extent of concordance between them in response to inductive signals. We demonstrated that while certain genes have a high degree of concordance of change between expression and accessibility changes, there is also a large group of differentially expressed genes whose local chromatin remains unchanged. By comparison, data from cell types along the hematopoietic differentiation trajectory had a much higher degree of concordance between genes with large gene expression changes and chromatin accessibility changes.

What might explain the lack of concordant changes in chromatin accessibility? One explanation could be that pre-existing chromatin accessibility dictates the *de novo* binding of transcription factors, but that the binding of transcription factors to those regions does not result in further changes to accessibility. Such effects have been reported in the context of glucocorticoid signaling, in which the glucocorticoid receptor almost exclusively binds to chromatin that is already accessible in response to dexamethasone [29]. Indeed, we demonstrated that genes that lacked concordance between changes in chromatin accessibility and gene expression were more likely to have nearby chromatin that was already accessible (Figures 3C,D). It is possible that in MCF-7 cells, the transcriptional effects of RA and TGF-β do not lead to a significant change in the activity of pioneer transcription factors, which are able to bind directly to condensed or inaccessible chromatin to facilitate its opening [30]. Also, implicit in our approach is the assumption that an increase in accessibility is associated with an increase in expression, which is not necessarily the case if a genomic locus becomes accessible to a repressive factor or a bound repressive factor is displaced by a nucleosome.

We looked at MCF-7 cells exposed to retinoic acid and TGF-β because these two signals induce a robust transcriptional response through distinct mechanisms. RAR-α remains bound to DNA and interacts with transcriptional activators in response to retinoic acid binding, while SMAD family members require TGF-β to bind to surface receptors to translocate to the nucleus. Yet, despite these differences, we observed that many genes changed expression independent of changes in chromatin accessibility for both signals. It is, however, possible that signaling molecules that exert their effects through very different types of transcription factors may have a different profile of concordance between changes in accessibility and gene expression. It is possible that other types of factors in a different context (e.g., different cell line) may yield a stronger correspondence.

Our data characterized molecular changes resulting from a single input (retinoic acid or TGF-β) in a clonal cell line, whereas the majority of work reporting a stronger concordance between simultaneous measurements of accessibility and transcription compared entirely different cell types or cells undergoing a directed differentiation protocol. What we have observed in the case of a single perturbation applied to cells that are not thought to change type *per se* is increased or decreased transcription with less concomitant nearby change in accessibility. How can one reconcile these observations? One possibility is that if we were to leave the signal on for longer, or combine it over time with the effects of several other signals, that we eventually would observe many further changes in accessibility proximal to a gene, concordant with the aforementioned results from comparisons between cell types. Whatever the source, these further changes in accessibility do not seem to occur randomly, given that they largely reflect the direction of change in transcription (increased accessibility for upregulation, decreased for downregulation). It may be that these subsequent changes in accessibility do not explicitly change transcription, but rather alter the underlying regulatory logic of the gene; i.e., the removal of a signal may not lead to a decrease in the gene’s transcription, or the gene’s transcription may be sensitized or desensitized to some other set of transcription factors.

## Methods

### PCA of RNA and ATAC-sequencing samples

Principal component analysis and visualization of RNA-seq and ATAC-seq samples was performed using raw counts and performing a variance stabilizing transform. Results were visualized using functions from the R DESeq2 package [31].

### RNA-sequencing analysis

Initial RNA sequencing analysis was performed as previously [32]. Briefly, reads were aligned to the hg38 assembly using STAR v.2.7.1a and counted uniquely mapped reads with HTSeq v0.6.1 and hg38 GTF file from Ensembl (release 90). We used DESeq2 v1.22.2 in R 3.5.1 using a minimum absolute-value log-fold-change of 0.5 and a q value of 0.05. For genes with multiple annotated transcriptional start sites, we used the ‘canonical’ transcription start site from the knownCanonical table from GENCODE v29 in the UCSC Table Browser.

We performed functional over-representation and gene set enrichment analysis [21] of upregulated transcripts in the high dose retinoic acid and high dose TGF-β using clusterProfiler v4.0.5 and enrichplot v1.12.3 [33]. P values for the over-representation analysis were adjusted using a false discovery rate approach. We used the C5 ontology and H hallmark curated gene sets from the Molecular Signatures Database (MSigDB) v7.4 [19,20] as reference gene sets to compare our upregulated genes to.

### ATAC-sequencing analysis

ATAC-seq alignment and peak calling was performed as previously [17]. We aligned peaks to the hg38 assembly using bowtie2 v2.3.4.1, and filtered out low-quality alignments with samtools v1.96, removed duplicate read pairs with picard 1.96, and used custom Python scripts along with bedtools v2.25.0 to create alignment files with inferred Tn5 insertion points. We called peaks using MACS2 [34] v2.1.1.20160309 with the command, ‘`macs2 callpeak --nomodel --nolambda --keep-dup all --call-summits -B --SPMR --format BED -q 0.05 --shift 75 --extsize 150`’.

Since we had three biological replicates per condition, we used a majority rule approach to retain only summits that were found in at least two replicates [35]. Using these condition-specific peak files, we used bedtools to create a consensus peak file by merging each individual condition’s peak summit file together in a manner that disallowed overlapping peaks. We used bedtools merge command ‘`bedtools merge -d 50`’ to combine features within 50 base pairs of each other into a single peak after testing multiple merge distances. We used the number of ATAC-seq fragment counts at each peak in this merged consensus peak file for differential peak analysis.

We used the custom peak analysis algorithm from Sanford et al., 2020 that took advantage of additional ethanol control conditions to estimate false discovery rate in ethanol controls to then identify differential peaks. Briefly, reads were quantified for each peak in the master consensus file and fragments at each peak were normalized to correct for differences in total sequencing depth using the equation: sample’s total reads in peaksmean number of reads in peaks across all samples. Next, an estimated false discovery rate was calculated in each cell of a 50×50 grid containing 50 exponentially-spaced steps of minimum fold-change values (ranging from 1.5-10) and 50 exponentially-spaced steps of minimum number of normalized fragment counts in the condition with the greater number of counts (ranging from 30 to 237 or 10 to 237). The estimated false discovery rate (FDR) was calculated using the equation: estimated FDR = (no. of conditions)(est. number of false positive peaks per condition)total number of differential peaks in experimental conditions. After calculating the estimated FDR in each cell of the 50×50 grid, we then pooled together differential peaks contained in any cell with an FDR less than 0.25%.

We performed motif analysis on our set of differential peaks using chromVAR v1.8.0 [22], its associated cisBP database of transcription factor motifs, and the motifmatchR package from bioconductor. To decrease the variance of the transcription factor motif deviations scores, we pooled together the different dosages of retinoic acid or TGF-β. The chromVAR code was modified to extract an internal metric that equals the fractional change in fragment counts at motif-containing peaks for a given motif.

### Hematopoietic differentiation data

We used preexisting RNA- and DNase I-seq data (aligned to genome assembly hg19) of hematopoietic differentiation [11] to compare against our data. We used data from the website provided in the paper (http://cbio.mskcc.org/public/Leslie/Early_enhancer_establishment/) to download annotations of peaks (peaksTable.txt), counts of DNase-seq (DNaseCnts.txt), and RNA-seq counts (RNAseqCnts.txt). Counts presented in these data files were quantile normalized and averaged when biological replicates were available. We filtered peaks with “CD14” or “CD34” under the “accessPattern” annotation to choose for peaks that were relevant for comparing HSPCs to monocytes. We used a log_2_ fold change of greater than or equal to 2 as a cutoff for assigning differential peaks. We used the preexisting annotations of genes for each peak for peak-gene mapping. For determining the log_2_ fold change in gene expression we discarded genes whose maximum expression value across the two conditions was fewer than 5 quantile-normalized units.

For visualization of this data set with our own accessibility data, we realigned raw fastq files DNase-seq files to the hg38 assembly using bowtie v2.3.4.1 and filtered out low-quality alignments with samtools v1.1 to generate new .bam alignment files. The alignment files were combined using samtools merge in a single .bam file per cell type. Bam files were converted to .bigWig format using deeptools 3.5.1 [36] “`bamCoverage --normalizeUsing CPM`” to create a ‘consensus’ .bigWig for visualization. Peaks for CD34+ and CD14+ samples were made by filtering peaks annotated for these populations in the “accessPattern” column and creating separate .bed files using a custom script. The peak location in these .bed files were then lifted over from hg19 to hg38 using UCSC hgLiftOver. For comparing the overlap of peaks between data sets, we created consensus peak sets across all sample types and used the bedtools intersect function to quantify the proportion of peaks that intersected between the hematopoietic differentiation and signaling data.

### Peak annotation

Peaks were annotated using ChIPseeker [28] to determine the relative proportion of features in the data from González et al., 2015 (DNAse-seq) and Sanford et al., 2020 (ATAC-seq). For ease of visualization, certain categories like the three promoter categories were collapsed into one. ChiPseeker was also used to identify the nearest transcriptional start site to a gene used for the nearest integration approach described below. For making scatter plots of change in accessibility versus change in expression annotated by peak feature, a custom script was used to combine annotations from ChIPseeker into four categories: downstream, gene body, integenic, and promoter.

For each of the top 150 most variable transcription factor motifs we identified using differential accessibility analysis, we used the R bioconductor motifmatchR package to annotate both the number of motif matches and a log-likelihood match score for each peak.

### RNA and ATAC data integration

We employed two methods for assigning peaks to genes. In the ‘nearest’ approach, we used annotation from ChIPseeker to assign each peak to the nearest transcriptional start site. With this method, each peak is uniquely mapped to a single gene. In the ‘window’ approach we used a window of 50 kilobases on either side of the transcriptional start site (100 kilobases in total) to assign peaks to a gene, which could result in a peak being assigned to multiple genes.

### Track Visualization

We visualized accessibility data using the web based version of integrative genomics viewer (IGV) [37,38]. We prepared accessibility data for visualization by taking consensus files and converting them to .bigWig file format with either fragments per million or counts per million normalization. Bed files for identifying peaks were created using custom scripts.

### Statistics and software

Unless otherwise stated, we performed analyses using R v4.1.0 with data manipulation and visualization done with tidyverse v1.3.1 [39] and ggpubr v0.4.0. We used a Kolmogorov-Smirnov test to compare means. Unless otherwise stated, 95% confidence intervals for Pearson’s r were calculated by bootstrapping using 10,000 replicates.

## Data and Code Availability

All raw and processed data as well as code for the analyses in this manuscript can be found at: https://www.dropbox.com/sh/qbjuagz511c072g/AAChvYMjdoG7A0eNdqbEmaUla?dl=0

## Author Contributions

KK and AR conceived and designed the project with input from EMS and YG. KK designed and performed all analyses supervised by AR. KK performed data representation and visualization. EMS and YG contributed some input to analysis and data visualization. KK and AR wrote the manuscript with input from all authors.

## Acknowledgements

We are greatly indebted to Professor Christina Leslie and Alvaro González for many insightful discussions and for assistance in working with their datasets. We also thank the members of the Raj lab for valuable feedback, especially Ally Coté and Lee Richman. KK acknowledges support from NIH T32 GM008216; EMS acknowledges support from F30 HG010986; YG holds a Career Award at the Scientific Interface from BWF; and AR acknowledges support from NIH Director’s Transformative Research Award R01 GM137425, NIH R01 CA238237, NIH R01 CA232256, NIH P30 CA016520, NIH SPORE P50 CA174523, and NIH U01 CA227550.

