## Supplemental Figures for "Changes in chromatin accessibility are not concordant with transcriptional changes for single-factor perturbations"

**Supplemental Figure 1: Global analysis of expression and chromatin accessibility changes in response to varying signals in MCF-7 cells.**

a. PCA of variance stabilizing transformed raw counts from gene expression and chromatin accessibility data demonstrating the first two principal components.

b. Gene set enrichment analysis (GSEA) [21] of differentially expressed genes in response to high dose retinoic acid against a gene set for skeletal system morphogenesis. Genes whose expression were differentially expressed in response to TGF-β were enriched for genes associated with epithelial-to-mesenchymal transition. Green traces represent running enrichment scores across fold change ranked gene lists.

**Supplemental Figure 2: Tuning peak calling parameters.**

a. Representative peak calls at the *CYP26A1* using different peak merge parameters (colors) and minimum normalized fragment count coverage (shades of the same color). Based on these results we selected a merge distance of 50 base pairs and a minimum coverage of 30 normalized fragment counts.

**Supplemental Figure 3: Comparison of accessibility data from hematopoietic differentiation and MCF-7 cells in response to signal.**

a. Number of differentially expressed genes (left) specific to CD34+ hematopoietic stem and progenitor cells (HSPCs, blue) and CD14+ monocytes (orange) from data from González et al., 2015 and the number of differentially accessible peaks (DNase-seq) between the two populations (right).

b. Annotation of distribution of peak location in relation to gene transcriptional units for consensus files for HSPCs and monocytes (left). Distribution of accessible peak features for consensus peaks for MCF-7 cells in ethanol, high dose retinoic acid, and high dose TGF-β.

**Supplemental Figure 4: Comparison of peak calls at multiple loci across both hematopoietic differentiation and MCF-7 genome-wide accessibility data sets.**

a. Consensus peak calls for MCF-7 signal samples (ATAC-seq) and hematopoietic differentiation samples (DNase-seq) at a 'housekeeping' gene (*GAPDH*), hematopoietic cell-specific marker loci (*CD34, CD14*), and retinoic acid and TGF-β responsive sites (*DHRS3, SERPINA11*). Values are fragments per million for ATAC-seq samples and counts per million for DNase-seq samples.

**Supplemental Figure 5: Expression and peak width distributions in MCF-7 signal data based on locus complexity.**

a. $log_2$-transformed expression of low complexity (teal) and high complexity genes (green) in response to retinoic acid (left) and TGF-β (right). P-values represent the probability of these data or more extreme under the null hypothesis that the distribution of gene expression values were drawn from the same probability distribution via the Kolmogorov-Smirnov test.

b. Distribution of peak widths for low complexity (teal) and high complexity (green) peaks with the median peak width (151 base pairs) marked by the dotted black line.

**Supplemental Figure 6: Concordance between gene expression change and proportion of differentially accessible peaks per gene for high and low complexity genes using a lower minimum coverage threshold for differential peaks.**

**Supplemental Figure 8: Accessibility-concordant and accessibility-non-concordant genes have similar loci complexity and differences in peak accessibility after exposure to signal depending on change in gene expression.**

a. Distribution of loci complexity the four groups of genes with differential expression in response to high dose retinoic acid.

b. Distribution of loci complexity the four groups of genes with differential expression in response to high dose TGF-β.

c. Accessibility after exposure to high dose retinoic acid. Accessibility of every peak assigned using the 'nearest' approach for gene groups based on accessibility concordance. P-values represent the probability of these data or more extreme under the null hypothesis that the distribution of peak accessibilities were drawn from the same probability distribution via the Kolmogorov-Smirnov test.

d. Accessibility after exposure to high dose TGF-β. Accessibility of every peak assigned using the 'nearest' approach for gene groups based on accessibility concordance. P-values represent the probability of these data or more extreme under the null hypothesis that the distribution of peak accessibilities were drawn from the same probability distribution via the Kolmogorov-Smirnov test.

**Supplemental Figure 9: Effect of window size on number of differentially accessible peaks based on gene expression change and correlation of gene expression and accessibility changes using medium and low dose signals.**

a. Distributions of number of differentially accessible peaks for differentially expressed and non-differentially expressed genes in response to high dose retinoic acid (left) or high dose TGF-β (right) based on window size around transcriptional start site (TSS).

b. 'Nearest' approach to assigning peaks to genes shows less concordance in signaling compared to hematopoietic differentiation. Scatter plots showing change in peak accessibility (median or maximum) versus

$\log_2$ fold change in expression on y axis for medium and low dose retinoic acid (first two columns) and medium and low dose TGF-β (second two columns). Pearson's correlation coefficients reported with 95% confidence interval from bootstrapping with 10,000 replicates in parentheses.

c. 'Window' approach to assigning peaks to genes shows less concordance in signaling compared to hematopoietic differentiation. Scatter plots showing change in peak accessibility (median or maximum) versus $\log_2$ fold change in expression on y axis for medium and low dose retinoic acid (first two columns) and medium and low dose TGF-β (second two columns). Pearson's correlation coefficients reported with 95% confidence interval from bootstrapping with 10,000 replicates in parentheses.

#### Supplementary Figure 1.

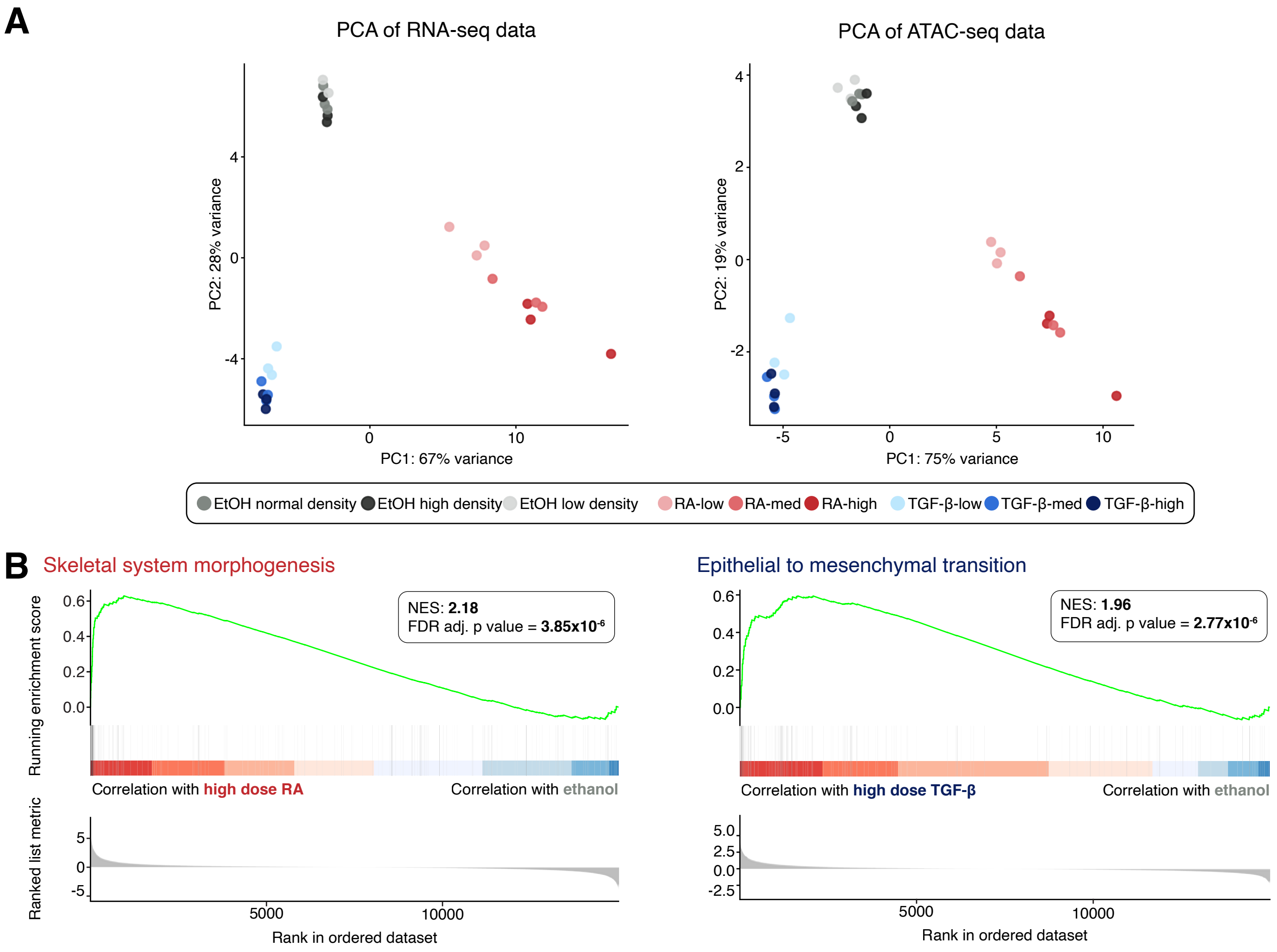

### Supplemental Figure 2.

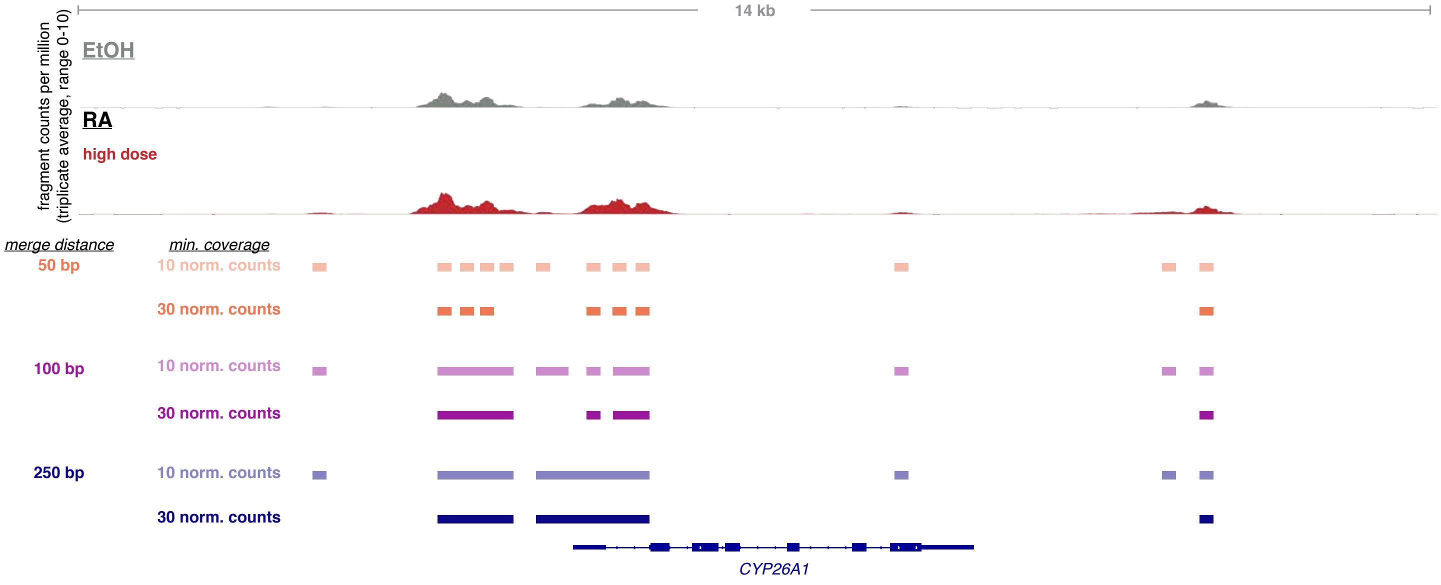

### Supplementary Figure 3.

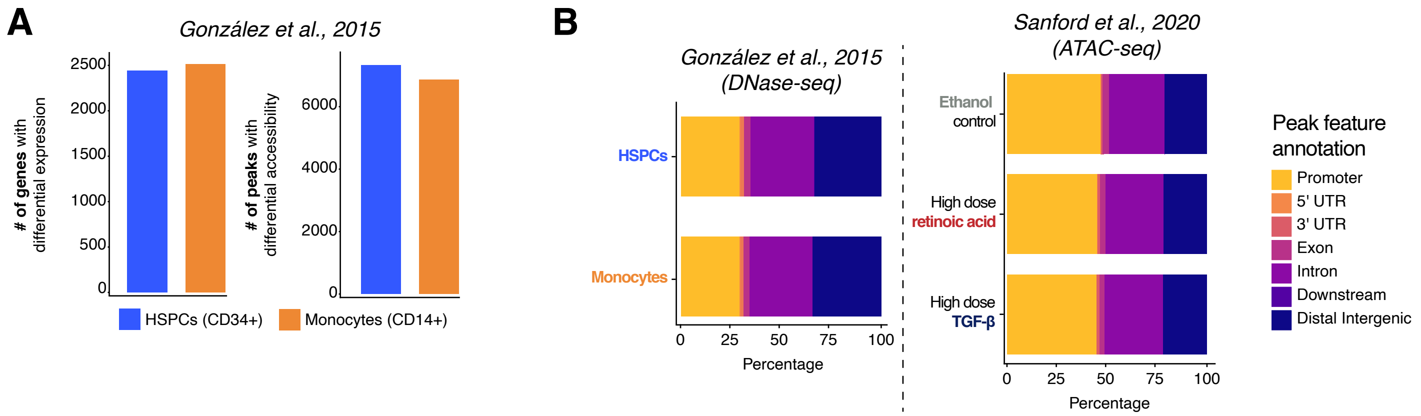

#### Supplemental Figure 4.

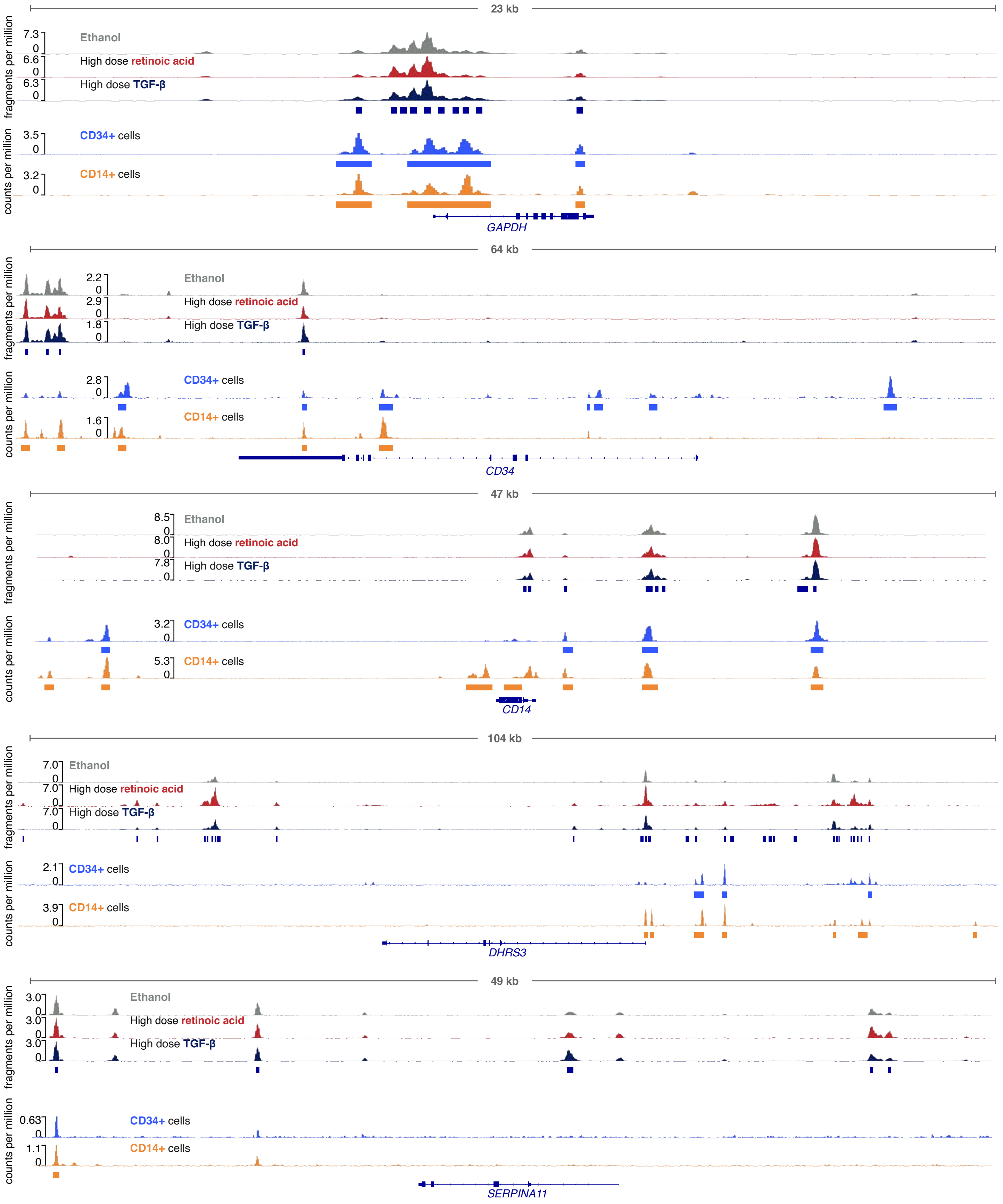

#### Supplemental Figure 5.

A

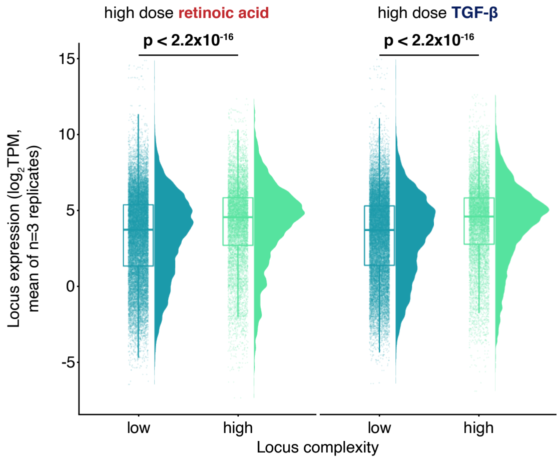


B

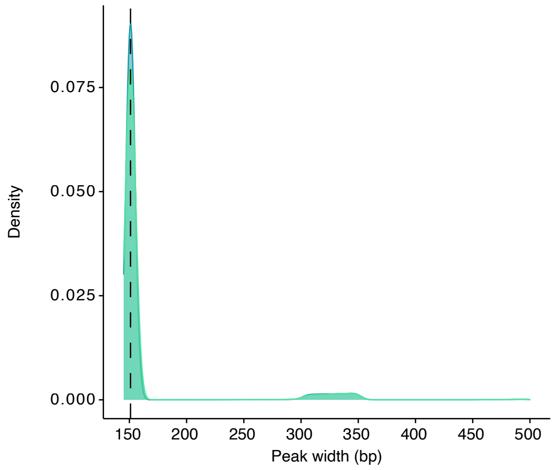

### Supplemental Figure 6.

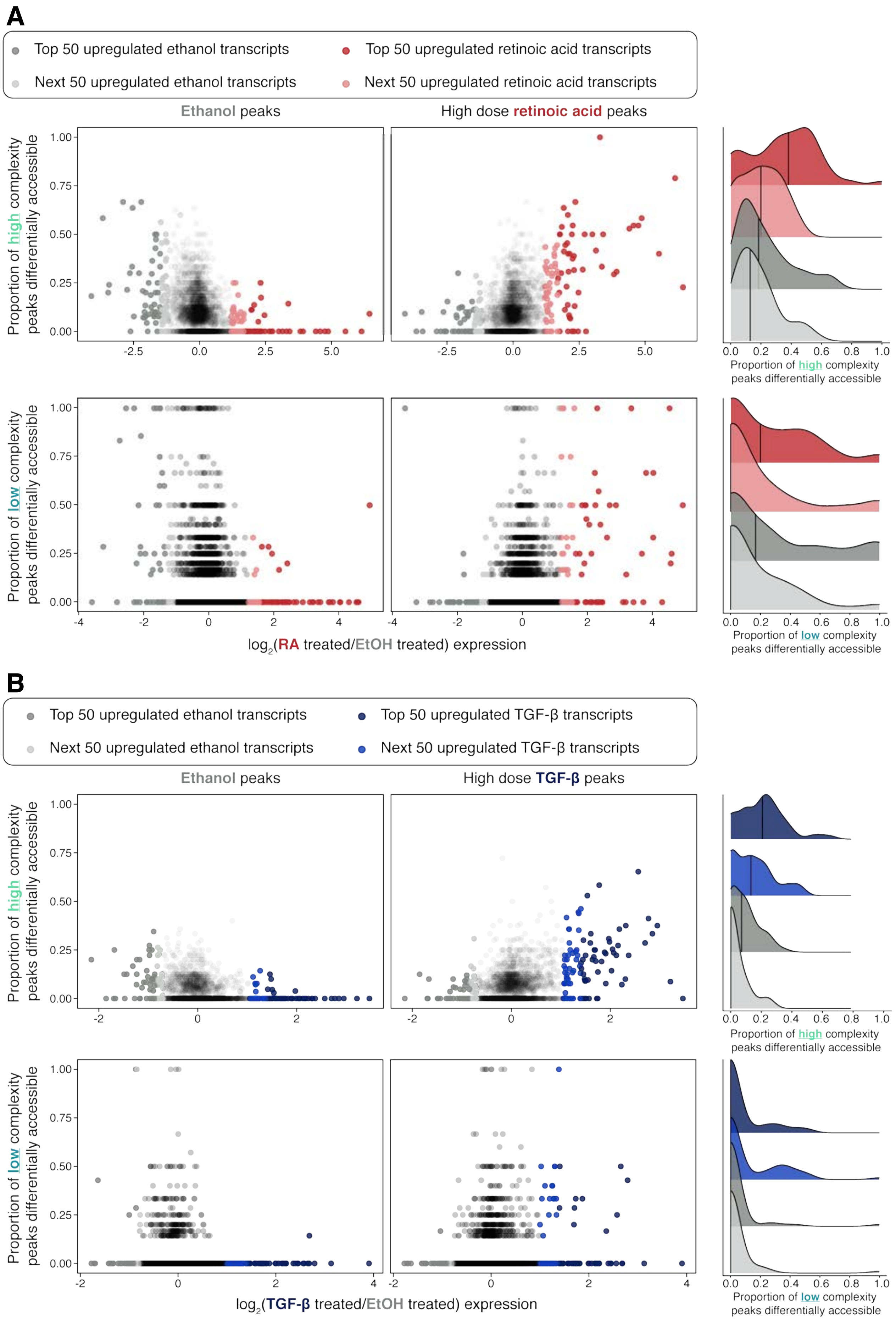

#### Supplemental Figure 7.

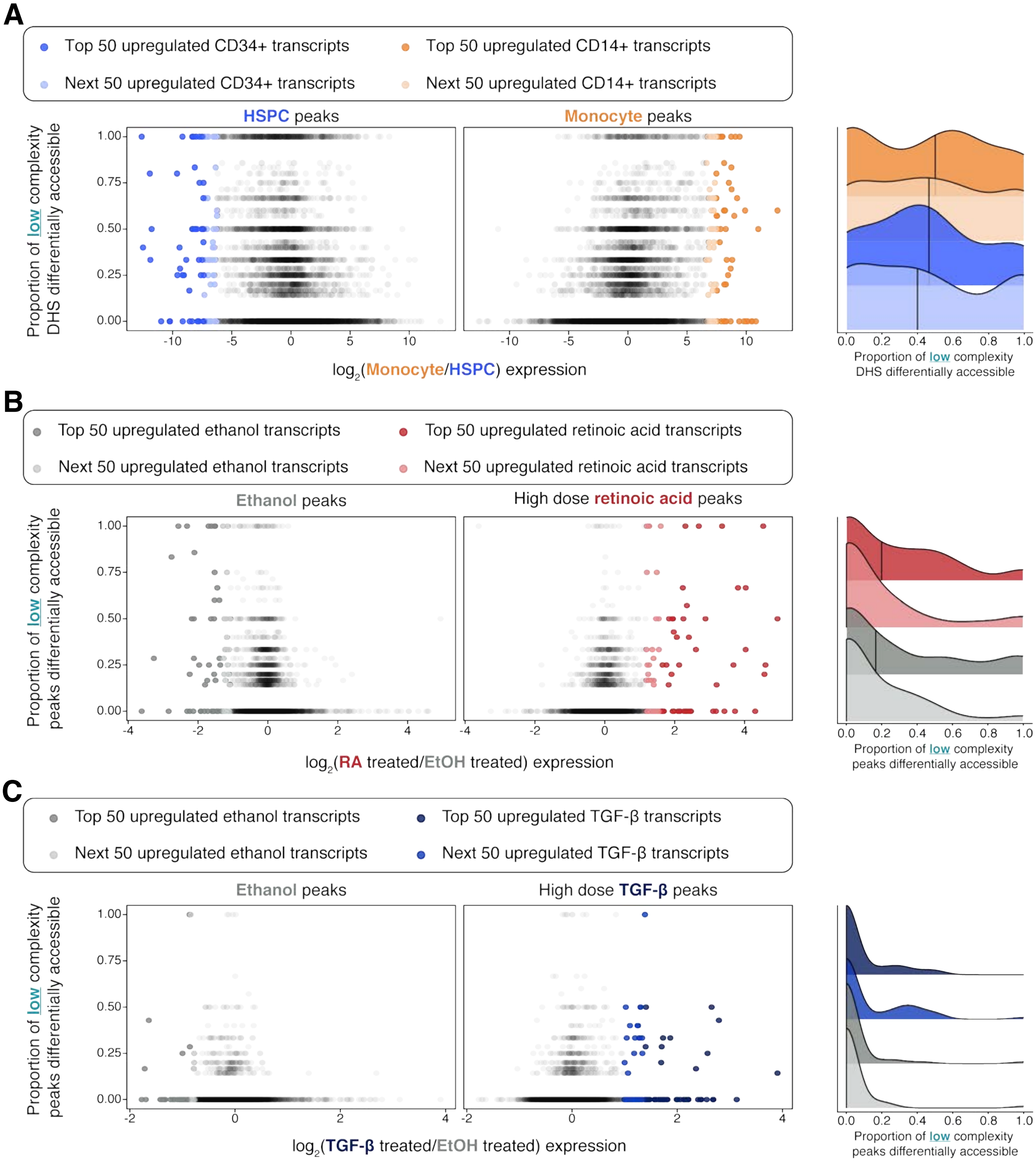

#### Supplemental Figure 8.

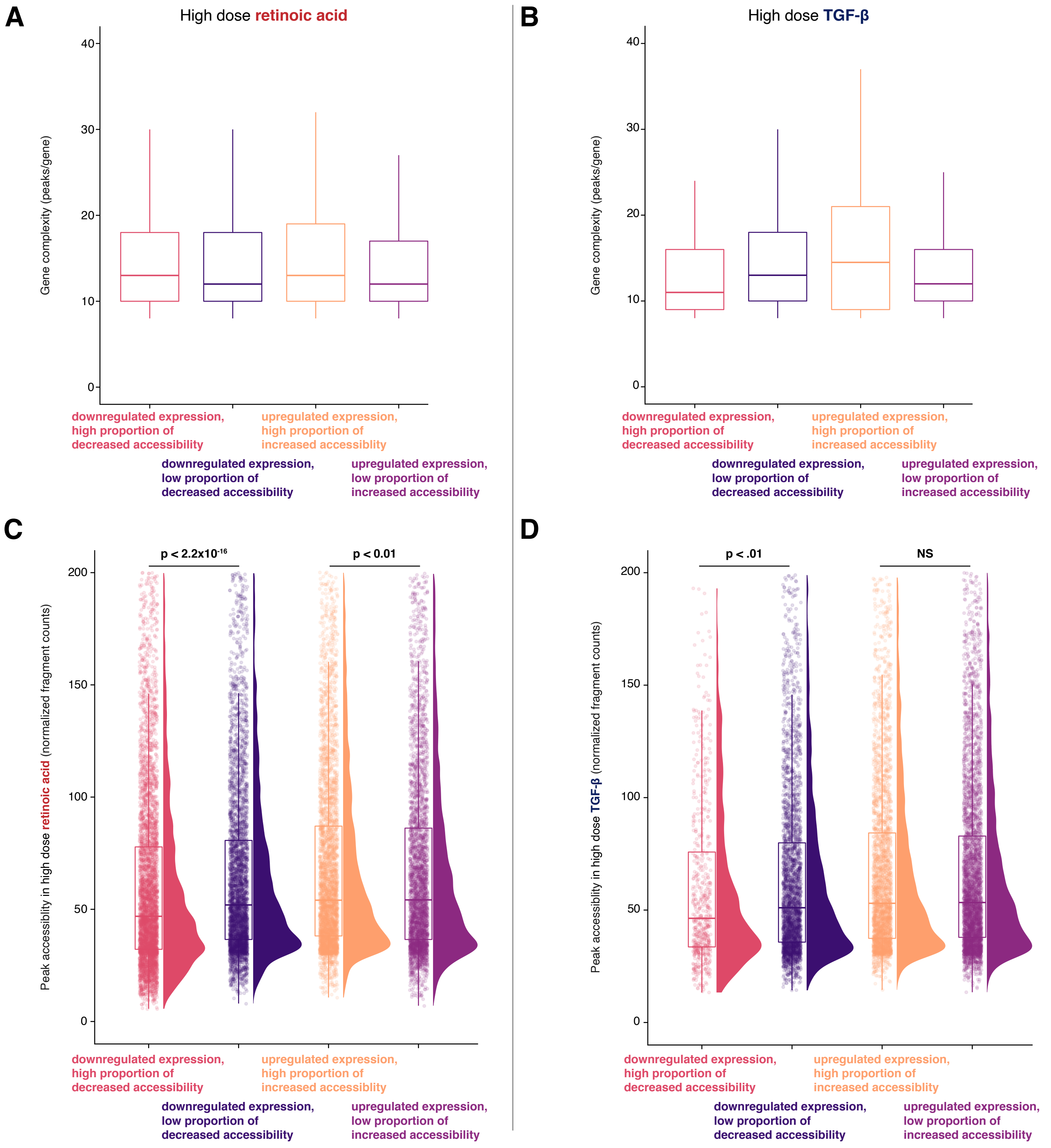

### Supplemental Figure 9.

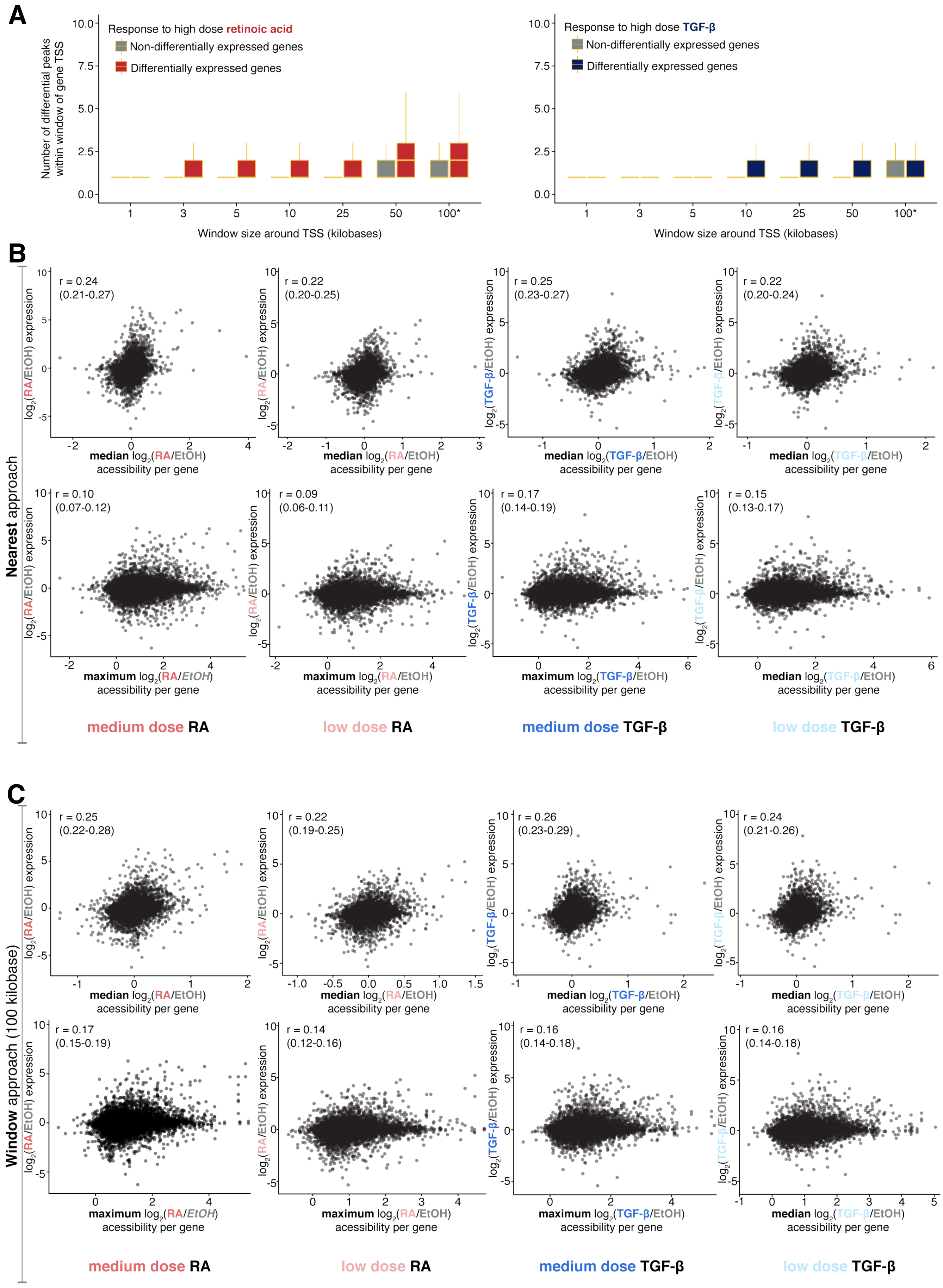
